# Context and exposure history shape insecticide sensitivity in *Drosophila melanogaster*

**DOI:** 10.64898/2026.09.28.753401

**Authors:** Andjela Plavotić, Bettina Ansorge, Emma Sunderbrink, Anne-Kathrin Classen

## Abstract

Insecticide toxicity is commonly assessed under standardized conditions, yet susceptibility may depend on the biological state of the organism, its environment, and its recent exposure history. How these factors shape susceptibility at the organismal level remains insufficiently understood. Here, using *Drosophila melanogaster*, we systematically compared responses to three insecticides with distinct modes of action - acetamiprid, chlorantraniliprole, and deltamethrin. Susceptibility varied substantially with developmental stage, sex, and age, while temperature altered toxicity in a compound-dependent manner, demonstrating that lethal effects are strongly conditioned by biological and environmental context. Susceptibility was also rapidly modified by prior exposure: sublethal pre-exposure to each insecticide reduced lethality upon subsequent challenge with the same compound, with increased tolerance emerging within 24 h. However, exposure conditions that enhanced insecticide tolerance were associated with reduced resistance to heat stress and nutrient starvation, indicating that increased tolerance to an insecticide challenge can coincide with reduced resilience to other environmental stresses. Sublethal exposure also affected reproductive performance: increasing acetamiprid concentrations reduced egg production but surprisingly increased developmental success of the resulting offspring, largely maintaining the number of offspring reaching adulthood. Together, our findings identify rapid, compound-specific, and potentially costly tolerance as an underappreciated consequence of insecticide exposure and establish insecticide susceptibility as a dynamic organismal trait shaped by physiology, environmental conditions, and recent exposure history. These findings highlight the importance of incorporating such sources of variation, together with the broader consequences of sublethal exposure, into pesticide risk assessment.

## INTRODUCTION

Insecticides are indispensable to modern agriculture, but their extensive use raises concern for non-target organisms and ecosystem health (Hassaan & El Nemr, 2020). Residues of current-use insecticides are routinely detected in soil, water, vegetation, and non-target insects well beyond treated fields (Brühl et al., 2021; Herrero-Hernández et al., 2020; Mauser et al., 2025). This is particularly concerning against the backdrop of well-documented insect decline, including the dramatic loss of flying insect biomass reported from protected areas in Germany (Hallmann et al., 2017). Research in different insect species, such as honeybees, cockroaches or grasshoppers, revealed that even sublethal exposure to insecticides can alter the insect physiology and fitness, which may contribute to insect decline (Albacete et al., 2024; Bantz et al., 2018; Charpentier et al., 2014; Desneux et al., 2007; F. Li et al., 2026). Beyond their ecological impact, repeated pesticide and insecticide exposure has also been linked to human health outcomes including cancer, endocrine disruption, and neurological disease (Pedersen & Hansen, 2026; Shekhar et al., 2024; Zhou et al., 2025).

*Drosophila melanogaster* offers an experimentally tractable system to dissect these effects from the organismal down to the molecular level. The functional genetic tractability and extensive molecular toolkit have made *Drosophila* a longstanding model for dissecting insecticide mode-of-action and or identify the genetic basis of pesticide resistance (Giansanti et al., 2025; Hayot et al., 2025; Holsopple et al., 2023; Homem et al., 2020; Kim et al., 2018; Rand et al., 2014; Troczka et al., 2015). Here, we use the quantitative power of the *Drosophila* system to systematically compare the organismal effects of three insecticides with distinct modes of action, namely acetamiprid, chlorantraniliprole and deltamethrin, chosen to span the major neuronal and neuromuscular target classes currently used in agriculture. Acetamiprid, a neonicotinoid, acts as an agonist of nicotinic acetylcholine receptors, thereby disrupting synaptic transmission and causing neuronal hyperexcitation, paralysis, and death (Bjørling-Poulsen et al., 2008; Hernandez-Jerez et al., 2024; Phogat et al., 2022). Chlorantraniliprole, an anthranilic diamide, activates insect ryanodine receptors, disrupting intracellular calcium homeostasis and resulting in uncontrolled calcium release, muscle contraction defects, paralysis, and death (Yang et al., 2023). Deltamethrin, a type II pyrethroid, targets voltage-gated sodium channels, prolonging channel opening and causing repetitive firing, paralysis, and death (Bjørling-Poulsen et al., 2008; Pitzer et al., 2021).

Using newly established pesticide exposure designs that allow for the reproducible delivery of very defined and sublethal insecticide concentrations (described in detail elsewhere), we show that in *Drosophila* sensitivity to all three compounds depends strongly on developmental stage, adult age, sex, and ambient temperature. To investigate whether short-term sublethal insecticide exposure modifies subsequent responses to insecticide challenge, we assess both the specificity of altered insecticide sensitivity and potential physiological trade-offs associated with prior exposure. We establish a quantitative organismal framework for comparing pesticide classes in *Drosophila* and define the phenotypic landscape onto which future molecular and mechanistic studies can be mapped, both to understand insecticide action in insects and, ultimately, to inform how a broader spectrum of pesticides may disrupt conserved biological processes relevant to human health.

## RESULTS

### Figure 1. Developmental stage and sex determine insecticide lethality in *Drosophila*

To compare how *Drosophila* larvae and adults responds to chemically distinct classes of insecticides, we focused on acetamiprid, chlorantraniliprole, and deltamethrin. Exposure assays were performed on a clear and preservative-free yeast extract/sucrose medium in glass vials. This optimized environment allowed us to compare dose-response relationships across developmental stages, sexes, and compounds under standardized, chemically inert and visually accessible conditions. We first performed dose-finding experiments to determine sublethal, lethal and LD50 concentration (causing 50% lethality) after 24 h of exposure in third-instar larvae, and in male and female adults starting at 3 days after eclosion. We defined organismal lethality by visible phenotypes: loss of mouth hook contractions in larval stages, and complete loss of appendage movement after vial tapping in adult flies. All assays (unless otherwise stated) were conducted at 18°C, chosen to improve insecticide chemical stability and reduce medium evaporation; the correspondingly slower development at this temperature also provided finer temporal resolution of post-exposure phenotypes and reduced inter-individual variability.

**Fig 1.**
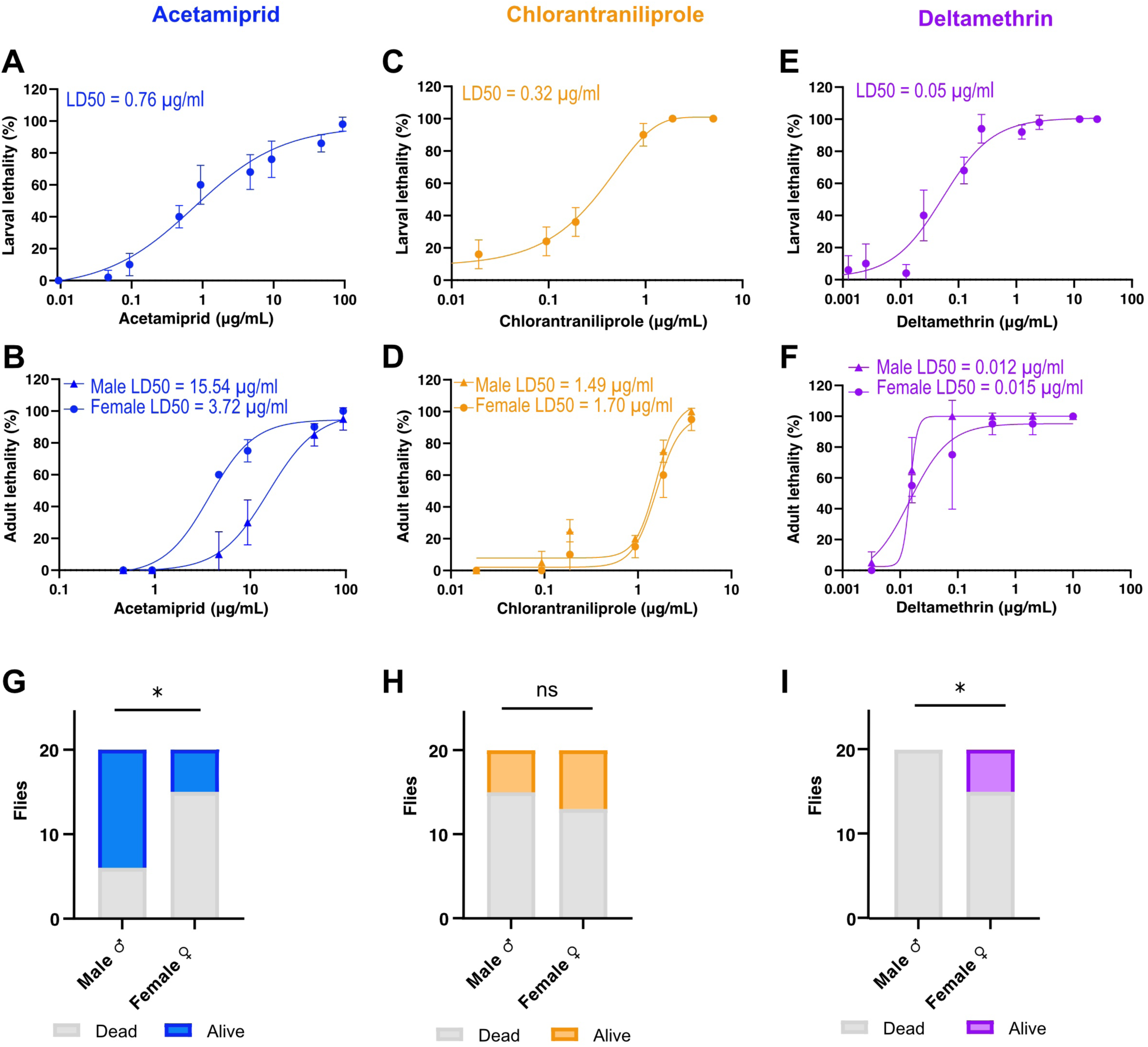
Dose-response relationships for three insecticides in larvae and adult male and female flies. **(A-F)** Dose-response curves for acetamiprid, chlorantraniliprole, and deltamethrin in larvae (A,C,E) and in adult male and female flies (B,D,F). Analysed using nonlinear regression fit model with variable slope. For larvae, n = 5 independent replicate vials of 10 larvae each were analyzed per concentration across 11 concentrations (N=550); for adult flies, n = 2 independent replicate vials of 10 flies each were analyzed per concentration across 7 concentrations (N=140). **(G-I)** Sex-specific responses to each insecticide in adults. Flies were exposed to acetamiprid (9.375 μg/mL), chlorantraniliprole (1.87 μg/mL), or deltamethrin (0.08 μg/mL), representing LD50 values obtain in October 2025. Statistical significance was assessed using Fisher’s exact test.

LD50 values differed markedly between developmental stages and sexes for all three compounds (**Fig 1A-F**). Fitted dose-response curves showed consistently lower LD50 values in larvae than in adults, identifying developmental stage as a determinant of susceptibility. Among adults, sex-specific sensitivity was compound-dependent: males were more sensitive than females to acetamiprid, females were more sensitive than males to deltamethrin, and no consistent sex bias was detected for chlorantraniliprole across independent experiments that used a range of sublethal to lethal concentrations (**Fig 1G-I, Fig S1A-C**). Together, these data show that insecticide sensitivity in *Drosophila* depends strongly on developmental stage, sex, and compound identity, rather than any one factor alone. In our hands, deltamethrin was particularly potent. In contrast to acetamiprid and chlorantraniliprole, we observed lethality for deltamethrin at concentrations that were over 10,000-fold lower than field application rates approved in the European Union (EU Commission, 2024). These results demonstrate that using defined exposure conditions in the *Drosophila* model provides a sensitive and highly quantitative system for analyzing biological responses to pesticide toxicity.

Over the course of our study, we observed seasonal variation in LD50 values despite tightly controlled laboratory conditions (**Fig S1D**). Similar seasonal variation in insecticide susceptibility has been reported, with higher susceptibility in autumn and lower susceptibility in summer and winter, patterns that may be influenced in part by environmental factors such as humidity (Miyo et al., 2000; Tillman et al., 2002). To account for this variability, we included matched control groups in all experiments. Unless stated otherwise, we performed the following LD50 exposure assays using a LD50 concentration determined in October 2025, which are slightly different form dose response curves shown in Fig 1 obtained in May 2026.

### Figure 2. Adult age and ambient temperatures modulate insecticide toxicity

Having determined that insecticide sensitivity differs by developmental stage and sex in *Drosophila*, we next asked whether additional physiological and environmental variables modify insecticide susceptibility. We focused on variables with direct ecological relevance: adult age, ambient temperature and duration of exposure.

**Fig 2.**
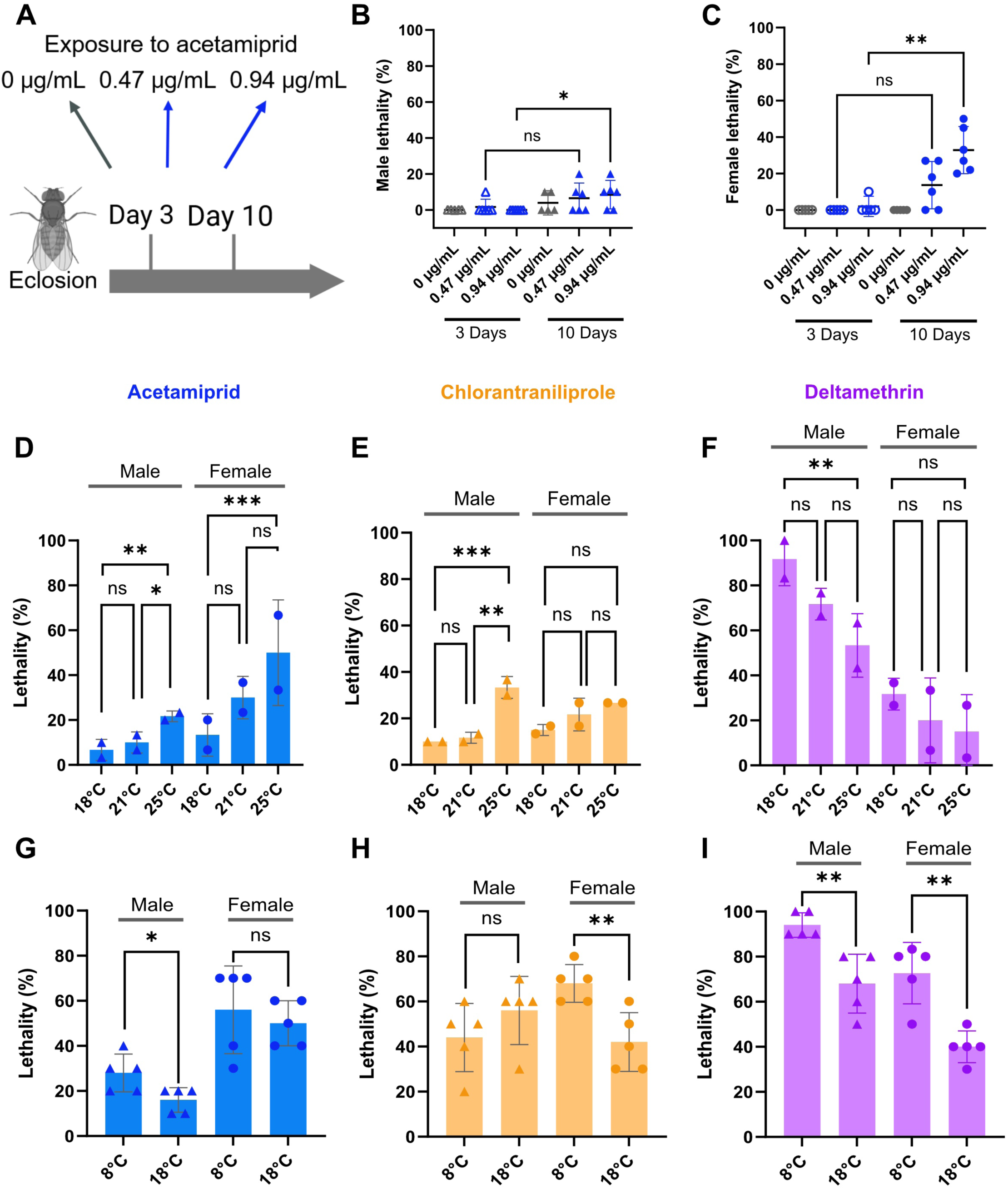
Impact of age and temperature on insecticide-induced lethality. **(A)** Schematic overview of exposure assay for adult flies on day 3 and day 10 post-eclosion. **(B-C)** Comparison of lethality between 3-day-old and 10-day-old male (B) and female (C) flies in control vials or exposed to two different acetamiprid concentrations. n = 5 vials with 10 flies each (N=50 flies) for controls and n = 6 vials for treatment groups (N=58-63 flies per condition). **(D-F)** Temperature-dependent lethality in response to an LD50 challenge of acetamiprid (D, blue, 5.5 μg/mL for male, 18.7 μg/mL for female), chlorantraniliprole (E, orange, 0.52 μg/mL for male, 1.73 μg/mL for female), and deltamethrin (F, magenta, 0.12 μg/mL for male, 0.017 μg/mL for female) exposure in male and female flies reared at 18°C, 21°C, and 25°C. Data was analyzed by a two-way ANOVA with Tukey’s multiple comparisons test. Two independent experiments are shown, where each experiment has n = 3 vials with 10 flies each (N=60 flies per condition). **(G–I)** Impact of cold stress on insecticide-induced lethality in response to the same LD50 concentrations as in (D-F) of acetamiprid (G, blue), chlorantraniliprole (H, orange), and deltamethrin (I, magenta) exposure in male and female flies reared at 18°C or 8°C. Data was analyzed with Welch’s t-test or the Mann-Whitney test as indicated by data distribution. n = 5 vials with 10 flies each (N=50 flies per condition).

We first tested whether adult age affects lethality after 24 h of acetamiprid exposure by comparing 3-day-old and 10-day-old flies. We found that 10-day-old flies were significantly more sensitive than 3-day-old flies, an observation reproducible in both sexes (**Fig 2A-C**). Thus, adult age even early in life represents an internal determinant of insecticide-induced lethality in *Drosophila*.

We next asked whether ambient temperature modifies insecticide-induced lethality, a question of direct ecological relevance given the substantial temperature variation insects experience in natural habitats, and the known influence of temperature on both insect physiology and toxicant efficacy. We first compared lethality at 18°C, 21°C and 25°C following exposure to acetamiprid, chlorantraniliprole, or deltamethrin. We found that acetamiprid- and chlorantraniliprole-induced lethality increased between 18°C and 25°C. In contrast, deltamethrin-induced lethality decreased between 18°C and 25°C (**Fig 2D-F, Fig S2**). To test if, in contrast to increasing temperatures, cold stress could also modify lethality, we compared insecticide-induced lethality at 18°C and 8°C (**Fig S2**). Housing flies at 8°C for 24 h alone caused no lethality, but strongly increased lethality induced by acetamiprid and chlorantraniliprole. For deltamethrin, low temperature further amplified the trend we already observed from 25°C to 18°C, with male lethality reaching over 90% at 8°C (**Fig 2G-I, Fig S2**). Thus, temperature modulated lethal outcomes in a compound-specific manner, with positive, negative, or neutral temperature coefficients readily observable.

### Figure 3. Sublethal exposure to insecticides induces insecticide-specific tolerance

In adult *Drosophila*, we observed visible behavioral phenotypes upon insecticide exposure. Acute abnormalities - tremor, shaking, abnormal posture, and impaired movement - appeared within seconds of exposure, even at low LD10 insecticide concentrations (**Supplement Video 1,2**). However, especially acute tremors appeared to decline over the first 24 h of exposure (**Supplement Video 3,4**). To better resolve how exposure duration shapes these visible responses, we defined four quantifiable phenotypic categories spanning the spectrum of pre-lethal severity: ‘dead’ (no movement after vial tapping), ‘supine’ (lying on back with visible leg movement), ‘ataxic’ (standing upright with minimal, uncoordinated leg movement), ‘hypoactive’ (standing upright with slow but coordinated walking). The cumulative distribution of these categories changed between 2 h and 24 h - driven by the increasing lethality to an LD50 exposure, but surprisingly, not between 24 h and 48 h, even when fresh acetamiprid was added at the 24 h timepoint (**Fig 3A-C**). These findings reveal that most lethality occurs within the first 24 h of an LD50 exposure which is then followed by a relatively stable phenotypic state, potentially consistent with an adaptive change in sensitivity to acetamiprid.

**Fig 3.**
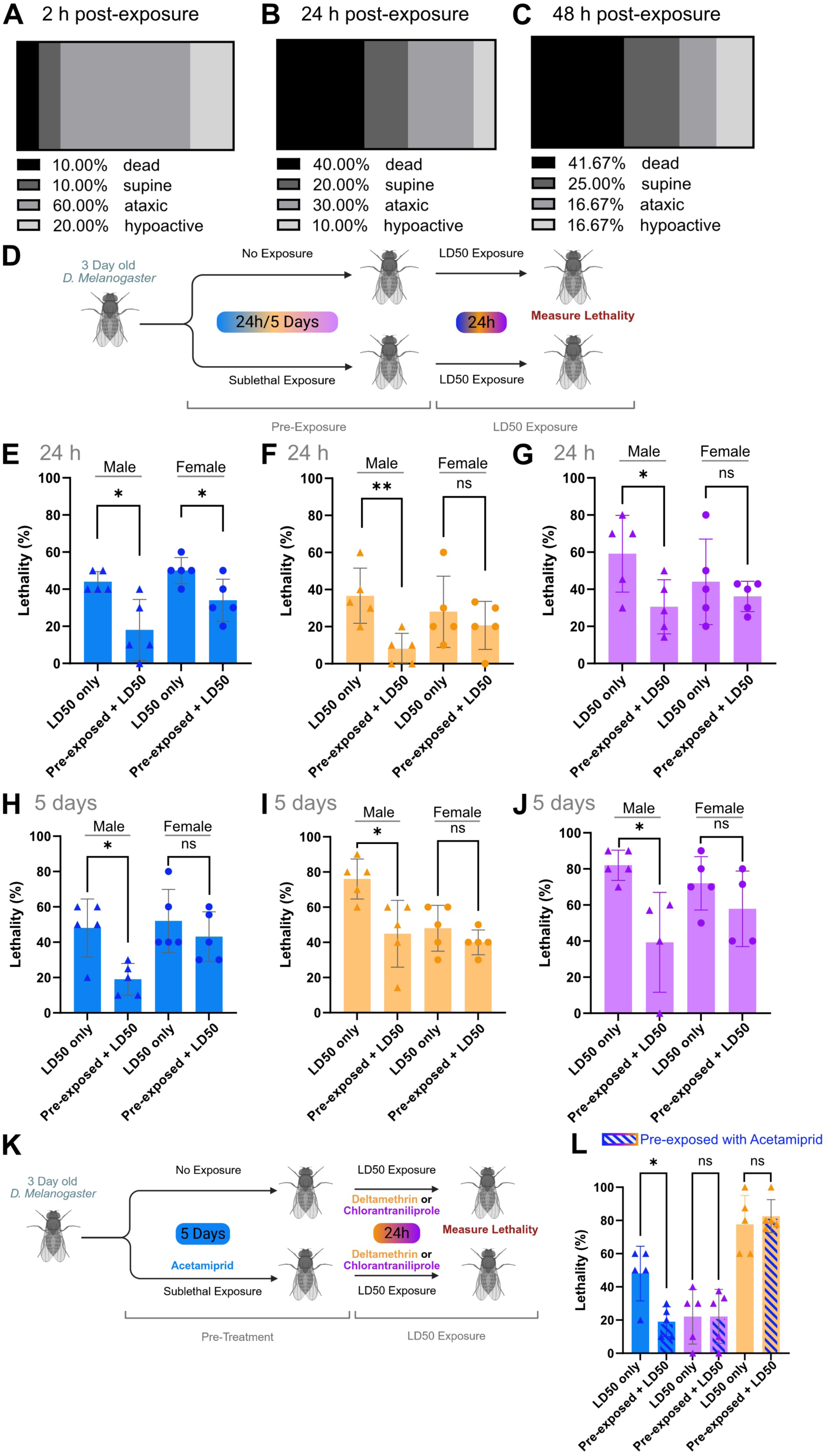
Insecticide tolerance emerges after sublethal pre-treatment. **(A-C)** Percentages of flies in different phenotypic states - ‘dead’ (no movement after vial tapping), ‘supine’ (lying on back with visible leg movement), ‘ataxic’ (standing upright with minimal, uncoordinated leg movement), ‘hypoactive’ (standing upright with slow but coordinated walking) - at 2 h, 24 h, and 48 h post-exposure to acetamiprid (9.38 μg/mL). n=9 vials with 10 flies each (N=90 flies). **(D)** Schematic assay workflow for sublethal pre-exposure followed by an LD50 challenge. **(E–G)** Lethality in response to a LD50 challenge following a 24 h control period or a 24 h period of sublethal pre-exposure with acetamiprid (E, 0.94 µg/mL, blue), chlorantraniliprole (F, 0.187 µg/mL, orange), and deltamethrin (G, 0.016 µg/mL, magenta). n = 5 vials with 10 flies each (N=50 flies) per condition. **(H-J)** Lethality in response to a LD50 challenge following a 5-day control period or a 5-day period of sublethal pre-exposure with acetamiprid (H, 0.94 µg/mL, blue), chlorantraniliprole (I, 0.187 µg/mL, orange), and deltamethrin (J, 0.016 µg/mL, magenta). n = 5 vials with 10 flies each (N=50 flies) per condition. **(K)** Schematic assay workflow for sublethal pre-exposure to acetamiprid followed by an LD50 challenge with chlorantraniliprole or deltamethrin. **(L)** Lethality of male flies in response to a LD50 challenge with acetamiprid (blue), chlorantraniliprole (orange) or deltamethrin (magenta) with or without a 5-day period of sublethal pre-exposure with acetamiprid (0.94 µg/mL, blue stripes). n = 5 vials with 10 flies each (N=50 flies) per condition. LD50 concentrations used in this figure are for acetamiprid 12.8 µg/mL, for chlorantraniliprole 1.13 µg/mL and for deltamethrin 0.07 µg/mL. All data in this figure were assessed for normality using the Shapiro-Wilk test and analyzed with Welch’s t-test or the Mann-Whitney test as appropriate.

To examine this adaptive change more systematically, we asked whether a prior sublethal exposure to acetamiprid, chlorantraniliprole, or deltamethrin modifies the response to a subsequent LD50 challenge, using lethality as a simple, robust endpoint. We reasoned that even low insecticide concentrations, which affect neuronal function but would not cause lethality, might already be sufficient to induce physiological adaptation and even tolerance. To test this, we first developed sublethal conditions that we could apply to both a 24 h period of acute and a 5 day period of chronic insecticide exposure. We identified acetamiprid and chlorantraniliprole concentrations, where only approximately 10% of flies die after 5 days of exposure (**Fig S3**). The steep slope of the dose response curve and high toxic potency of deltamethrin makes it difficult to predict lethality of the same dose in subsequent experiments; however, our final conditions caused about a third of the flies to die after 5 days of treatment (**Fig S3**).

Importantly, when flies were pre-exposed to either acetamiprid, chlorantraniliprole or deltamethrin using sub-lethal concentration for either just 24 h or for the prolonged period of 5 days, we found that this pre-exposure strongly reduced lethality in response to a subsequent LD50 challenge (**Fig 3D-J**). These findings demonstrate that all three insecticides induce an acquired physiological tolerance to the insecticide on very short timescale within the lifetime of the insect.

To understand if acquired tolerance reflects a compound-specific mechanism or the activation of a general detoxification program, we tested whether sublethal exposure to acetamiprid also protected against an LD50 challenge by the other two insecticides. We found that a 5-day sublethal pre-exposure to acetamiprid did not confer cross-resistance to the other two insecticides (**Fig 3K,L**), indicating that acquired tolerance is specific to the inducing compound rather than reflecting a shared protective program. Together, these experiments demonstrate that sublethal exposure history can strongly reshape subsequent toxicological outcomes within hours to days; a timescale compatible with rapid phenotypic plasticity.

### Figure 4. Sublethal insecticide exposure reduces organismal resilience to other environmental challenges

Genetically encoded insecticide resistance, acquired through mutation and selection across generations, may come with a fitness cost: resistance-associated changes in target genes or detoxification loci can disrupt processes that would otherwise support growth, reproduction, and survival in the absence of insecticide (Freeman et al., 2021; Gul et al., 2023; Homem et al., 2020; Miyo et al., 2000). Whether the same trade-off may apply to an acquired physiological tolerance is less established.

**Fig 4.**
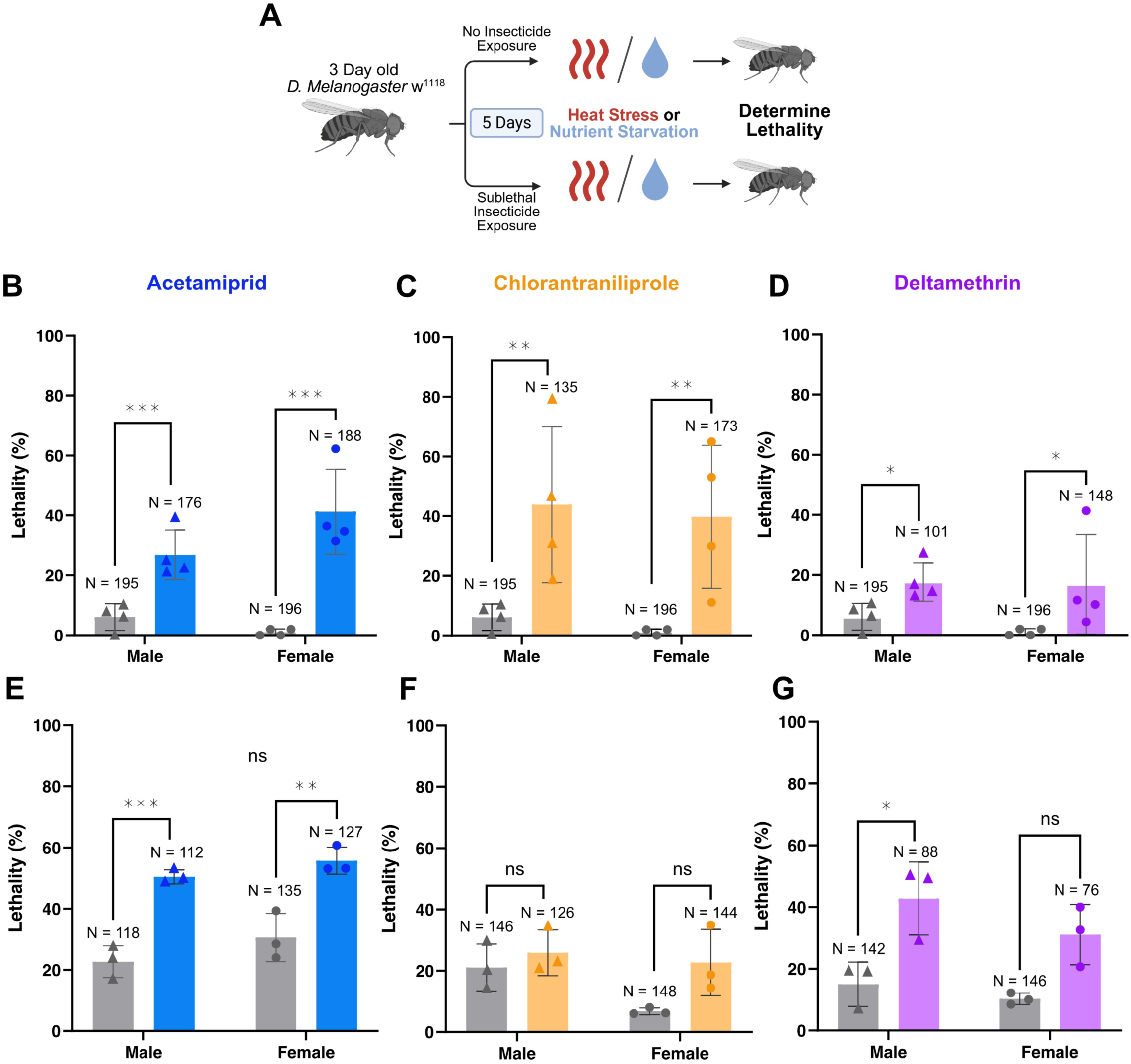
Stress resilience is reduced after prolonged sublethal insecticide exposure. **(A)** Schematic assay workflow to assess resistance to heat stress or nutrient starvation after a 5-day period of sublethal insecticide exposure. **(B-G)** Lethality upon exposure to 24 h of heat stress (32°C, B-D) or upon exposure to 66 h of nutrient starvation (on water only, E-G) following a 5-day period of sublethal insecticide exposure to acetamiprid (blue, 0.94 µg/mL, B, E), chlorantraniliprole (orange, 0.187 µg/mL, C, F), deltamethrin (magenta, 0.016 µg/mL, D, G). For heat stress resistance (B-D): Four independent experiments are shown, where each experiment has n = 5 vials with 10 flies each (N=200 flies per condition); total number of flies surviving sublethal pre-exposure for 5 days which were then exposed to heat stress were N = 195 or 196 for untreated controls and N = 101 - 188 for pre-exposed groups. For starvation resistance (E-G): Three independent experiments are shown, where each experiment has n = 5 vials with 10 flies each (N=150 flies per condition); total number of flies surviving sublethal insecticide pre-exposure for 5 days which were then exposed to starvation were N = 118-148 for untreated controls and N = 76-144 for pre-exposed groups. Data were analyzed by two-way ANOVA with Tukey’s multiple comparisons test.

We therefore asked whether flies that acquired physiological tolerance during sublethal insecticide exposure also showed deficits in fitness-related traits by focusing on two resilience read-outs: resistance to heat stress and resistance to nutrient starvation. To assess heat-stress resistance, we pre-exposed flies to sublethal concentrations of acetamiprid, chlorantraniliprole, or deltamethrin for 5 days at 18°C, then shifted them to 32°C onto insecticide-free medium and scored lethality after 24 h. Flies exposed to sublethal insecticide concentrations displayed increased lethality when shifted to 32°C if compared to untreated control flies (**Fig 4A-D**). To assess resistance to nutrient starvation, we transferred flies exposed to sublethal concentrations of acetamiprid, chlorantraniliprole, or deltamethrin for 5 days to vials containing only water and scored lethality after 66 h. Specifically flies pre-exposed to acetamiprid and deltamethrin displayed a significant increased lethality in response to nutrient starvation if compared with untreated, starved controls (**Fig 4A, G-E**). Together, these results show that the same 5-day sublethal exposure that conferred acquired tolerance to insecticide also reduced the ability of flies to withstand other, ecologically relevant stressors.

### Figure 5. Acetamiprid exposure alters reproductive outcome

To capture additional dimensions of organismal fitness after exposure, and specifically reproductive success, we assessed rates of egg laying and of developmental transitions all the way to adult eclosion. For this analysis, we exposed adult male and female flies to a low sublethal or a high LD50 concentration of acetamiprid for 5 days, while allowing them to mate. We then paired individual surviving females with two surviving males and measured egg laying over the following 4 days (**Fig 5A**). After low sublethal acetamiprid exposure, we found that egg-laying rates were mildly increased, but only within the first 24 h post-exposure, with no difference to controls on subsequent days (**Fig 5B**). Such a transient increase is consistent with hormetic responses reported for other sublethal xenobiotic exposures in insects reflecting a female-specific adaptive mechanisms increasing egg production under environmental stress (Cutler, 2013; Huangfu et al., 2024; Mandal et al., 2026). In contrast, females surviving 5 days of high LD50 exposure showed a chronically reduced egg-laying rate, which likely reflects how high acetamiprid exposure stress compromises the metabolically demanding process of egg production (**Fig 5B**) (Jiang et al., 2023; A. Zhang et al., 2022). Importantly, we observed that egg-hatching rates across all conditions were similar directly on day 1 of egg lay analysis, demonstrating that mating and fertilization during both low and high acetamiprid exposure conditions was successful (**Fig 5C**). However, surprisingly, rates of larval survival and pupariation were elevated in both high and low acetamiprid treated offspring (**Fig 5D**). Ultimately, high acetamiprid treated offspring thus strongly increased the total rate of developmental success from egg lay to adult eclosion if compared to controls. As a result, negative effects from the observed reduced egg output were balanced out, such that total fertility, i.e. the net number of offspring reaching adulthood, was comparable to controls (**Fig S4**). This observation may point to an adaptive process where developmental success and especially larval survival is promoted in offspring after environmental stress in female mothers, a conclusion supported by another recent study in *Drosophilidae* (Deans & Hutchison, 2022).

**Fig 5.**
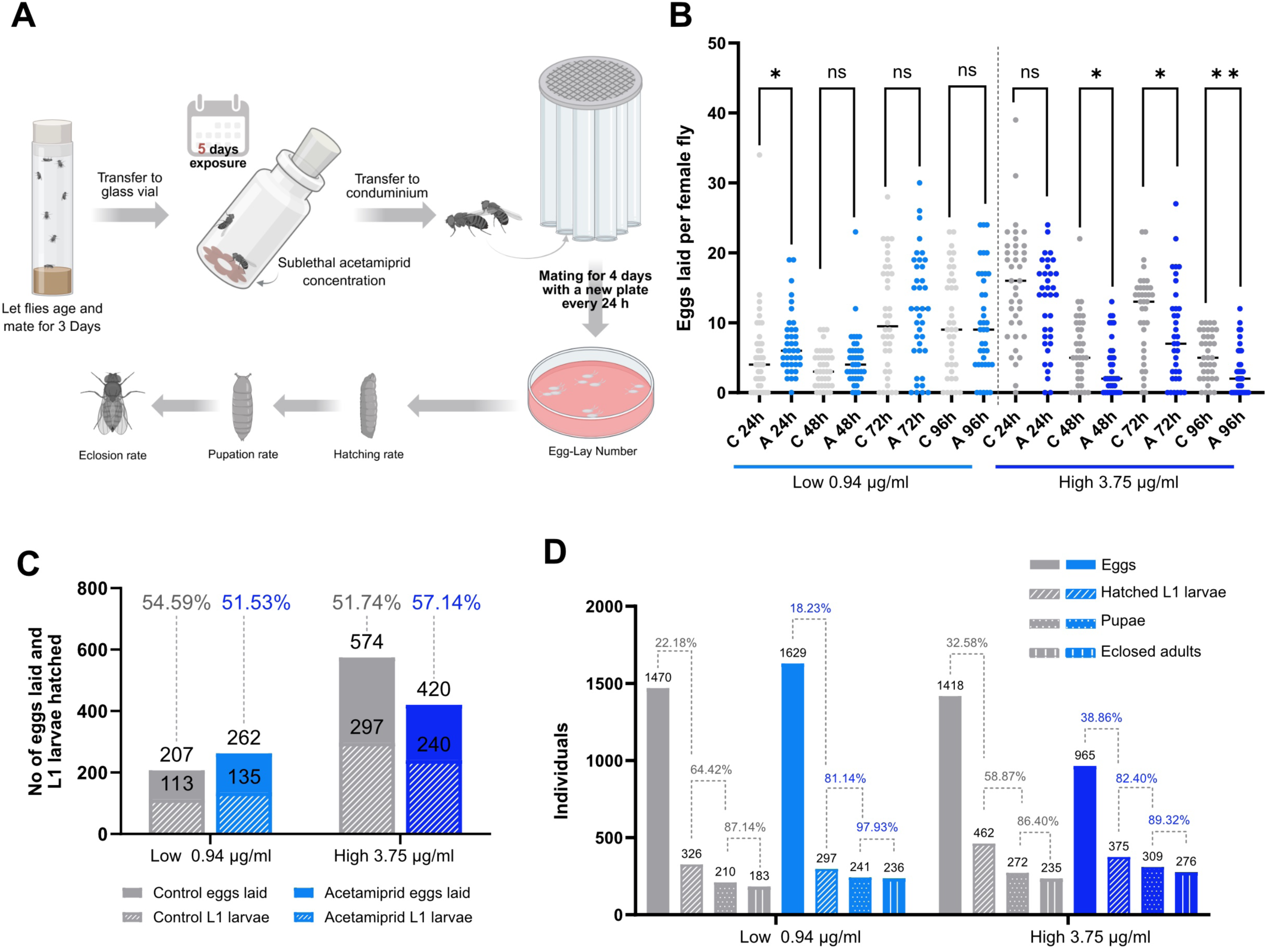
Sublethal acetamiprid exposure impacts fecundity and fertility. **(A)** Schematic workflow for the fecundity and fertility assay following a 5-day acetamiprid exposure. **(B)** Egg-laying rates after a 5-day period of exposure to either a low (0.94 μg/mL) or a high (3.75 μg/mL) acetamiprid concentration. Eggs were counted every 24 h for a 4-day period. C = untreated controls, A = acetamiprid-treated flies. n=38 females, each paired with two males per condition. **(C)** Egg-hatching rate from eggs collected during the first 24-h period after low and high acetamiprid exposure. Similar hatching rates suggests similar mating success and fertilization rates during acetamiprid exposure. **(D)** Total number of eggs laid, larvae hatched, larvae pupariated and adults eclosed. Numbers indicate percentage of individuals successfully transitioning to the next developmental stage.

## DISCUSSION

Using a quantitative *Drosophila* exposure system, we find that susceptibiliy to insecticides is strongly dependent on both internal and external variables. Sensitivity to acetamiprid, chlorantraniliprole, and deltamethrin varied with developmental stage, sex, adult age, and ambient temperature, extending observations previously made for individual compounds and insect species (Kalmouni et al., 2025; Miyo et al., 2000; Miyo & Oguma, 2002; Sedlmeier et al., 2025; Tasman et al., 2021). Larvae were consistently more sensitive than adults, whereas sex differences depended on insecticide identity. Although differences in body size, physiology, or detoxification capacity have been proposed to explain sex-dependent insecticide sensitivity in other insects (Sedlmeier et al., 2025), the opposing sex biases observed for acetamiprid and deltamethrin argue against a simple relationship between body size and susceptibility in *Drosophila*. Adult age represented another important source of variation, with a few days older flies showing increased sensitivity to acetamiprid. Increased insecticide susceptibility with advancing adult age has also been observed in *Drosophila* exposed to other insecticides (Colinet et al., 2016; Smirle et al., 2017; Sur et al., 2018).

Temperature likewise strongly modified insecticide lethality, but its effect depended on the compound we tested. Acetamiprid and chlorantraniliprole were generally more lethal at higher temperatures, whereas deltamethrin showed the opposite relationship. Increased toxicity of acetamiprid and chlorantraniliprole at elevated temperatures has also been reported in other insects (Abbes et al., 2015; Gandara et al., 2024; Kumar et al., 2023; Y. Li et al., 2020; Ma et al., 2012). In contrast, the increased toxicity of deltamethrin at lower temperatures is consistent with the negative temperature coefficient previously described for pyrethroids in vitro and in vivo (Azhar & Khan, 2020; Chinn & Narahashi, 1989; Glunt et al., 2018; Hinks, 1985; Khan & Akram, 2014; Mansoor et al., 2015; Raj Boina et al., 2009; Salgado et al., 1989). A central finding of this study is that recent exposure history also changes subsequent insecticide sensitivity in a manner consistent with acquired or induced tolerance. Rapidly induced pesticide tolerance has previously been demonstrated in other organisms, most extensively in amphibians, where early sublethal exposure can alter sensitivity to later pesticide challenge (DiGiacopo & Hua, 2020; Hua et al., 2014). Our results extend this principle to three mechanistically distinct insecticides in *Drosophila* and demonstrate that exposure history can modify toxicological outcomes over a short timescale.

The mechanism underlying this acquired tolerance remains unresolved. The rapid onset of the phenotype is compatible with physiological plasticity, including altered expression or activity of insecticide targets, transporters, or detoxification pathways. Insects can transcriptionally regulate cytochrome P450s, glutathione-S-transferases, esterases, and insecticide target proteins following xenobiotic exposure, and individual members of these families can show considerable substrate specificity (Bantz et al., 2018, 2022; Casida, 2011; Chen et al., 2019; Gao et al., 2026; Hafeez et al., 2020; Hua et al., 2014, 2015; Le Goff et al., 2006; Misra et al., 2011; Pfannenstiel et al., 2024; Sun et al., 2018; Willoughby et al., 2006; Witwicka et al., 2025; C. Wu et al., 2021; P. Wu et al., 2023; Yuan et al., 2026), which could potentially promote acquired tolerance. In our experiments, however, pre-exposure to acetamiprid did not increase survival during subsequent challenge to either chlorantraniliprole or deltamethrin. Thus, the tolerance induced by acetamiprid does not appear to result from a broadly protective response sufficient to protect against two other chemically and mechanistically distinct insecticides. Instead, it may involve compound-selective changes in xenobiotic metabolism or transport, or altered sensitivity at the insecticide target, mechanisms that can show pronounced compound specificity in *Drosophila* (Denecke et al., 2017; Le Goff et al., 2006; Perry et al., 2008). Establishing the mechanistic basis of the phenotype will require future molecular and pharmacological analysis.

The increased tolerance to challenge correlated with reduced resilience to other environmental stresses. Flies pre-exposed to sublethal insecticide concentrations showed increased sensitivity to heat stress, and pre-exposure to acetamiprid or deltamethrin also increased sensitivity to nutrient starvation. Genetically encoded insecticide resistance is frequently associated with trade-offs in other traits, although such costs depend strongly on the resistance mechanism and genetic background and are not universal (Freeman et al., 2021; Gul et al., 2023; Homem et al., 2020; Siddique et al., 2021; Tchouakui et al., 2020). Our results identify a conceptually related trade-off associated with short-term insecticide exposure: conditions that increase survival during a subsequent insecticide challenge can coincide with reduced capacity to withstand unrelated physiological stress. Whether the acquired tolerance itself causes these deficits, or whether tolerance and reduced stress resilience are parallel consequences of insecticide exposure, cannot yet be distinguished. Nevertheless, the results show that survival during a subsequent insecticide challenge alone does not capture the full physiological consequences of prior sublethal exposure.

Acetamiprid exposure also produced more complex effects on reproduction and offspring development. Low sublethal exposure caused a transient increase in egg laying during the first day after exposure, whereas females surviving the higher concentration showed persistently reduced egg laying. Biphasic reproductive responses to insecticides have been described in several insect species and are commonly discussed in the context of hormesis (Cutler, 2013; Huangfu et al., 2024; Mandal et al., 2026; Yu et al., 2010; Y. Zhang et al., 2021). In *Drosophila*, sublethal insecticide exposure can likewise stimulate reproductive performance (Deans & Hutchison, 2022), whereas stronger or chronic pesticide exposure can impair fecundity and ovarian function (Dissawa et al., 2025; Kishore et al., 2026). Unexpectedly, the reduced egg output following high acetamiprid exposure was accompanied by increased developmental success among the resulting offspring, particularly during larval development, such that the number of offspring reaching adulthood was not reduced relative to controls. The basis of the effect observed here remains unknown.

Taken together, our results show that insecticide toxicity in *Drosophila* emerges from an interaction between compound identity, physiological state, environmental conditions, and previous exposure. Developmental stage, sex, adult age, and temperature substantially altered susceptibility to LD50 doses, while a sublethal LD10 exposure rapidly changed the outcome of subsequent LD50 insecticide challenges. Importantly, this increased tolerance induced by sublethal exposure co-occurred with reduced resilience to other stresses, demonstrating that decreased sensitivity to a second insecticide exposure does not necessarily indicate an absence of physiological consequences. These findings establish *Drosophila* as a tractable system in which to dissect how short-term exposure history modifies insecticide susceptibility and to identify the molecular processes that connect insecticide tolerance with organismal physiology.

## ACKNOWLEDGEMENTS

We thank Andrew Straw, Jason Tennessen, Georg Petschenka, Carsten Brühl, Nicola Iovino and Katrin Kierdorf and all members of the lab for help, support and discussion during this project. We thank the IMPRS-EBM graduate school for supporting our doctoral researchers.

## AUTHOR CONTRIBUTIONS

Conceptualization AP, BA, AKC

Experimental Validation AP, BA, ES

Experimental Investigation AP, BA, ES

Writing AP, AKC

Visualization AP, AKC

Supervision AKC

Funding Acquisition AKC

## FUNDING

Funding for this work was provided by the Deutsche Forschungsgemeinschaft (DFG, German Research Foundation) under Germany’s Excellence Strategy (CIBSS - EXC-2189 - Project ID 390939984) and by the Boehringer Ingelheim Foundation (RiseUp! Programme) to AKC.

## MATERIALS AND METHODS

### Fly stocks and maintenance

*Drosophila* melanogaster flies (*w^1118^*) were maintained under standard laboratory conditions at 18 °C, 70 % relative humidity, and a 12:12 h cycle on standard fly food (10L water, 74.5g agar, 243g dry yeast, 580g corn flour, 552ml molasses, 20.7g Nipagin, 35ml propionic acid).

### Preparation of insecticide exposure medium and insecticide exposure assays

The insecticide exposure assays are extensively described for larval and adult stages elsewhere (protocols submitted). Briefly, both larval and adult food media were prepared without antibiotics, mold inhibitors, or other preservatives. For larval exposure assays, semi-solid food consisting of 100 mL water, 0.3 g agar, 4 g sucrose, and 5 g yeast extract was prepared. For adult exposure assays, liquid food containing 16% sucrose and 6% yeast extract in water was prepared and sterilized by autoclaving.

For larval exposure assays, insecticides were mixed directly into the semi-solid food before solidification while the medium was cooling during preparation, and 2 mL were transferred to a wide glass vial. 10 early third-instar larvae were added to each vial. For adult exposure assays, two daisy-shaped Whatman filter papers were placed at the bottom of a tall glass vial to retain the liquid food. Ten flies, aged to 3 days post-eclosion, were separated by sex under brief CO₂ anesthesia and transferred into each experimental vial. Flies were allowed to fully recover from anesthesia, as indicated by the resumption of normal movement, before 130 µL of liquid food medium was added through the vial plug using a syringe and needle.

### Insecticide exposure conditions and lethality assessment

Larvae and adult flies were maintained for 24 h on either insecticide-free control food or food containing acetamiprid (CAS 160430-64-8), chlorantraniliprole (CAS 500008-45-7), or deltamethrin (CAS 52918-63-5) dissolved in water. To establish dose-response-curves, larvae were exposed for 24 h, after which lethality was assessed based on the presence or absence of mouth-hook contractions. For adults, exposure was performed either for 24 h (acut) or for 5 days (prolonged, chronic). During prolonged exposure, flies were kept for only a maximum of 3 days in the same vial. On the third day, flies were transferred to a fresh glass vial containing new daisy-shaped Whatman filter paper, and 130 µL of fresh exposure medium. This transfer step reduced the risk of fungal or bacterial growth in the preservative-free food.

### Stress resilience assays after insecticide pre-exposure

Flies were exposed to sublethal concentrations of acetamiprid (0.94 µg/mL), chlorantraniliprole (0.187 µg/mL), or deltamethrin (0.016 µg/mL) for 5 days. After 3 days, flies were transferred to fresh vials containing freshly prepared exposure medium. Following the 5-day exposure period, flies were subjected to conditional stress. To assess heat stress resistance, flies were transferred to insecticide-free food and maintained at 32 °C for 24 h, after which lethality was quantified. To assess nutrient starvation resistance, flies were maintained on water as medium for only 66 h (empirically determined time point of longest viability in untreated controls), after which lethality was quantified.

### Fertility Assay

Flies were maintained under control or insecticide-exposed conditions for 5 days, with males and females housed together. After exposure, one female and two males were transferred to individual egg-laying condominiums containing grape juice agar supplemented with yeast paste. Flies were allowed to lay eggs for 24 h and were transferred to fresh plates with yeast paste daily for 4 consecutive days. Eggs were counted after each laying period, hatching was assessed 24 h later, and larvae were transferred to Nutri-Fly food to quantify pupation and adult eclosion.

---

**Fig S1.**
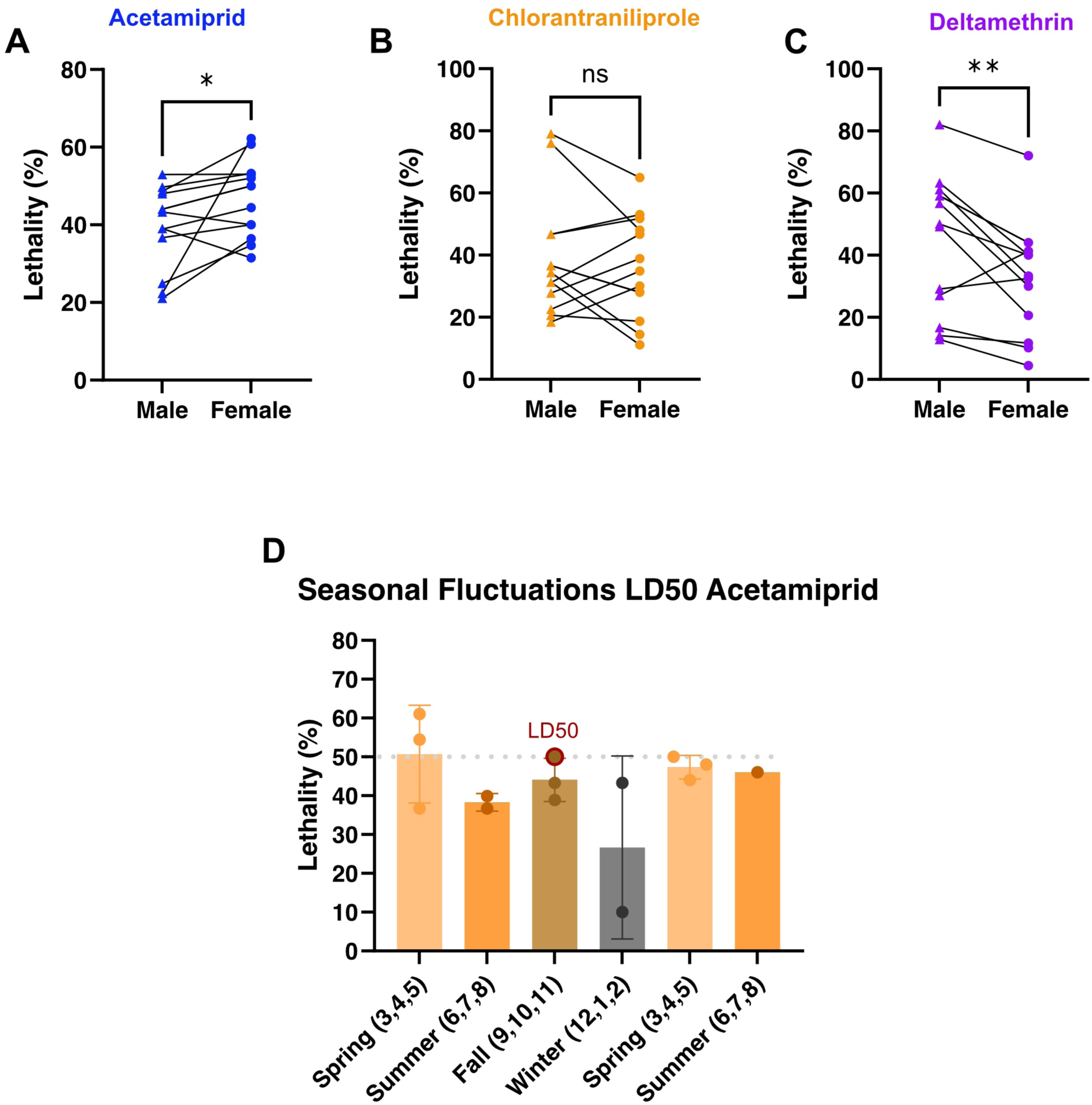
Sex specific responses and seasonal fluctuations in LD50 values. **(A-C)** Data from 13 independent experiments conducted over a 12-month period were pooled to assess sex-specific responses to acetamiprid (blue, 0.94 - 15.81 µg/mL), chlorantraniliprole (orange, 0.187 – 1.73 µg/mL), and deltamethrin (magenta, 0.016 – 0.11 µg/mL). Results are paired between the outcomes recorded for each sex in each experiment. Statistical significance was determined using a paired Wilcoxon signed-rank test. **(D)** Seasonal variation in acetamiprid-induced lethality. Flies exposed to the same acetamiprid concentration (15.54 μg/mL) during different seasons exhibit fluctuating lethality rates. Numbers in brackets represent the months of a year starting with 1 for January. Twelve individual experiments are shown, with n = 2-6 vials with 8-10 flies each per independent experiment.

**Fig S2.**
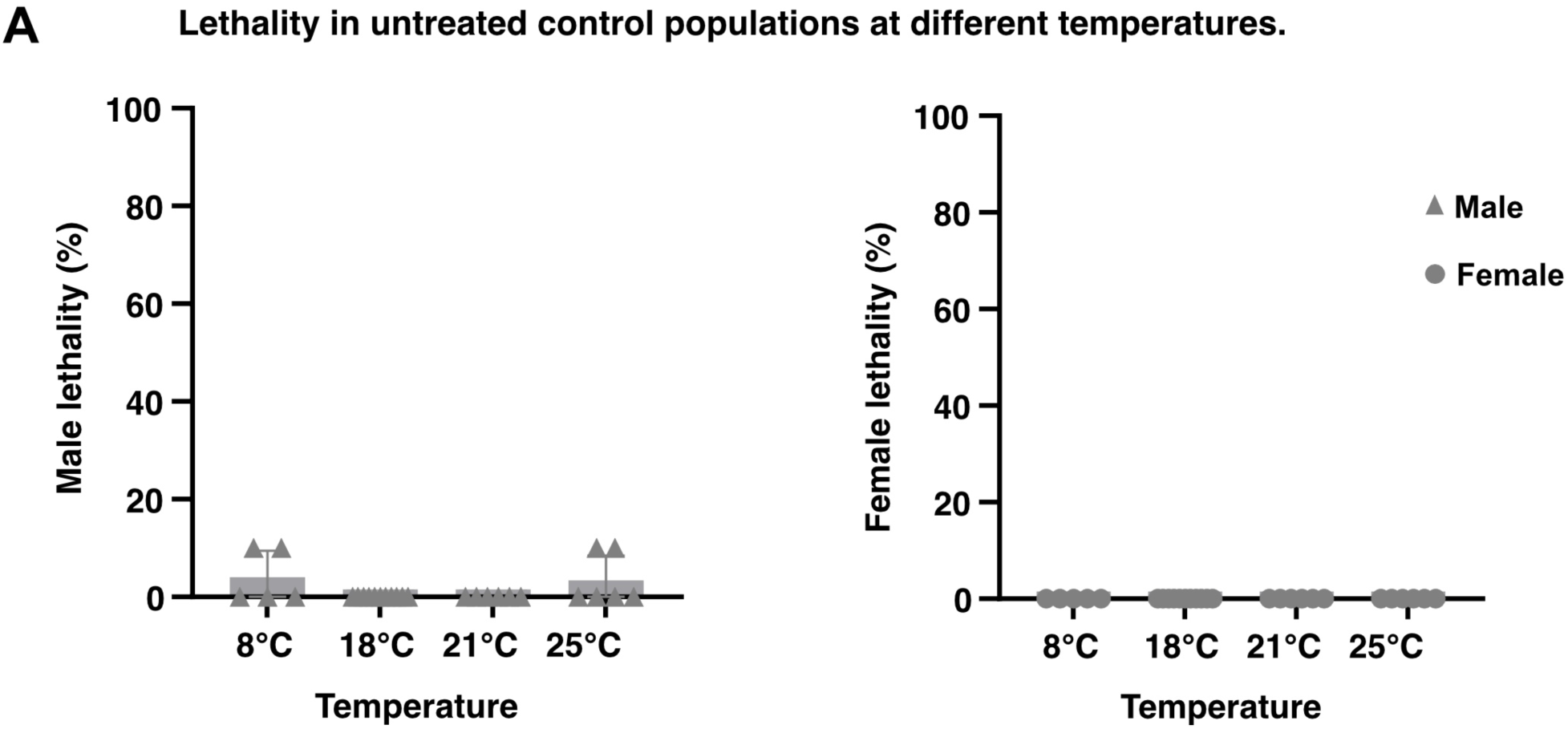
Lethality in control populations at different temperatures. **(A–B)** Male (A) and female (B) untreated control flies were maintained at four indicated temperatures and assessed for lethality after 24 h. For 8°C, 21°C, and 25°C, n = 5 vials with 10 flies per vial (N = 50 flies per temperature); for 18°C, n = 10 vials with 10 flies per vial (N = 100 flies).

**Fig S3.**
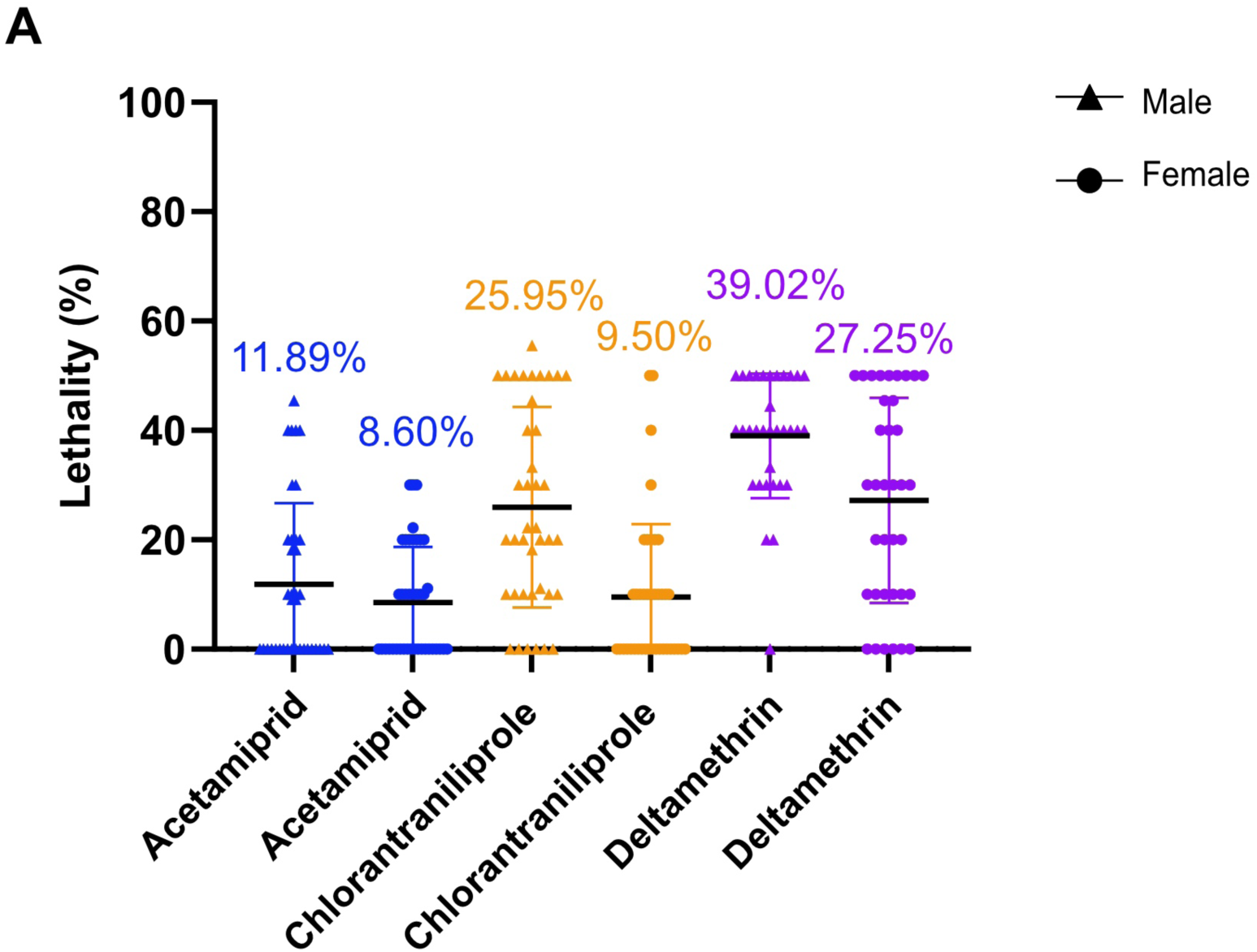
Survival rates of sublethal concentration exposure for 5 Days. **(A)** Lethality of male and female flies across multiple independent experiments in response to a 5-day period of sublethal exposure to acetamiprid (blue, 0.94 µg/mL), chlorantraniliprole (orange, 0.187 µg/mL), and deltamethrin (magenta, 0.016 µg/mL). Data were pooled from eight independent experiments performed separately for each insecticide and sex. For acetamiprid, the analysis included n = 40 vials comprising N = 397 female flies and n = 37 vials comprising N = 375 male flies. For chlorantraniliprole, n = 40 vials were analyzed for each sex, corresponding to N = 400 female flies and N = 396 male flies. For deltamethrin, the analysis included n = 36 vials comprising N = 362 female flies and n = 33 vials comprising N = 328 male flies.

**Figure S4.**
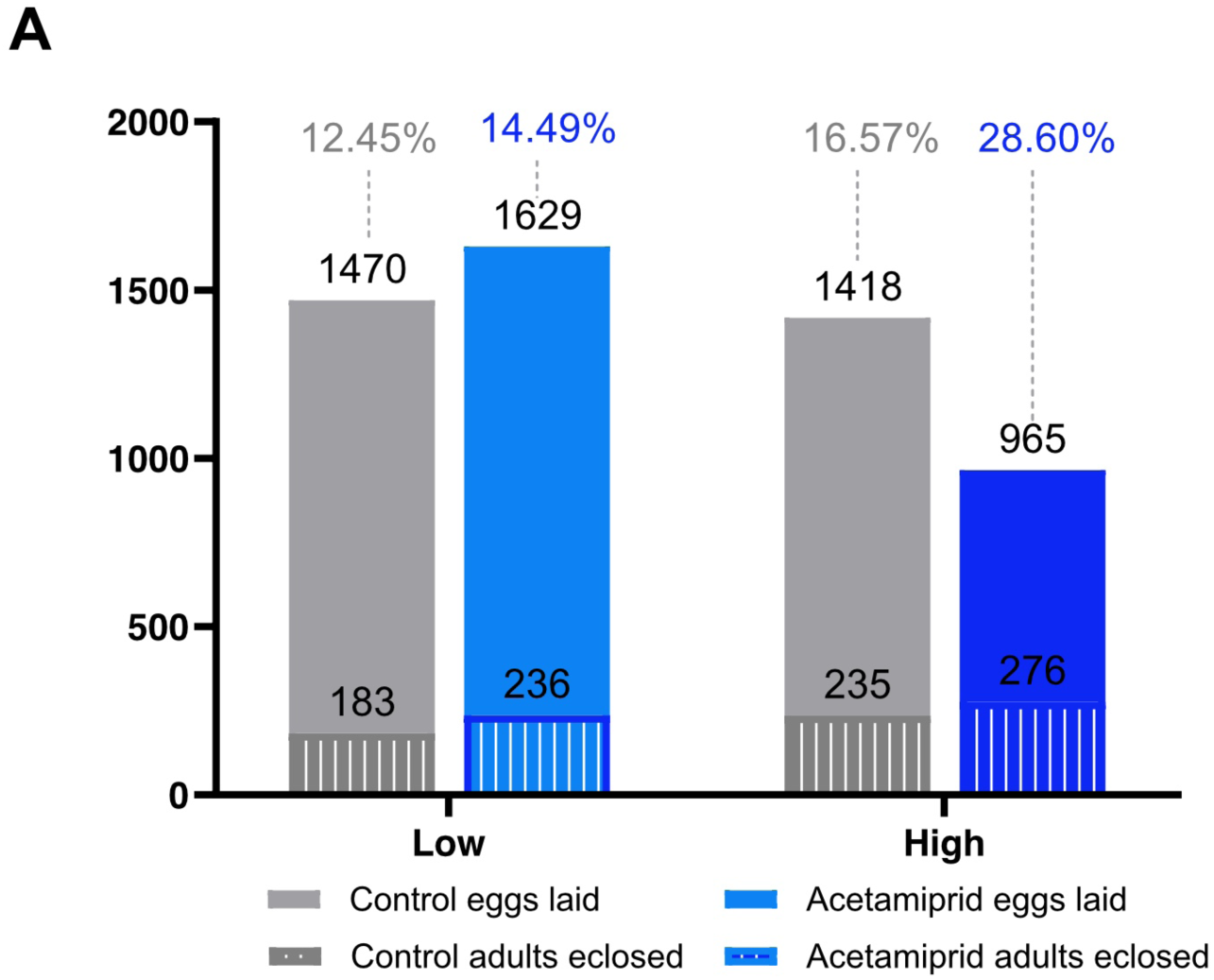
Total fertility rate following exposure to low and high concentrations of acetamiprid. **(A)** Total number of adults eclosed from the total number of eggs laid by females over a 4 day period after a 5-day untreated control period or after a 5-day period of exposure to either a low (0.94 μg/mL) or a high (3.75 μg/mL) acetamiprid concentration. Percentages represent final fertility rates for each condition.

**Supplementary Video 1. Response of adult flies after 5 min of sublethal insecticide exposure with mechanical stimulus.** Flies were exposed to sublethal concentrations of acetamiprid (0.94 µg/mL), chlorantraniliprole (0.187 µg/mL), or deltamethrin (0.016 µg/mL). The video was recorded 5 min after the onset of exposure and includes a single mechanical stimulus applied by tapping the vial to assess the flies’ mobility and behavioral response.

**Supplementary Video 2. Response of adult flies after 5 min of sublethal insecticide without stimulus.** Flies were exposed to sublethal concentrations of acetamiprid (0.94 µg/mL), chlorantraniliprole (0.187 µg/mL), or deltamethrin (0.016 µg/mL). The video was recorded 5 min after the start of exposure.

**Supplementary Video 3. Response of adult flies after 24 h of sublethal insecticide exposure with mechanical stimulus.** Flies were exposed to sublethal concentrations of acetamiprid (0.94 µg/mL), chlorantraniliprole (0.187 µg/mL), or deltamethrin (0.016 µg/mL). The video was recorded 24 h after the onset of exposure and includes a single mechanical stimulus applied by tapping the vial to assess the flies’ mobility and behavioral response.

**Supplementary Video 4. Response of adult flies after 24 h of sublethal insecticide without stimulus.** Flies were exposed to sublethal concentrations of acetamiprid (0.94 µg/mL), chlorantraniliprole (0.187 µg/mL), or deltamethrin (0.016 µg/mL). The video was recorded 24 h after the start of exposure.

## Notes

### Competing Interest Statement

The authors have declared no competing interest.

